# Translation components in adult *Drosophila melanogaster* adipocytes regulate the ovarian germline stem cell lineage

**DOI:** 10.1101/2024.08.31.610632

**Authors:** Subhshri Sahu, Alissa Richmond Armstrong

**Affiliations:** Western Carolina University, Department of Biology, Cullowhee, NC, USA; University of South Carolina, Biological Sciences, Columbia, SC, USA

**Author notes:** **Correspondence:** Alissa Richmond Armstrong.

**Keywords:** Drosophila, germline stem cells, translation, oogenesis, adipocytes

## Abstract

Adult stem cells, which support tissue homeostasis and damage repair, are influenced by whole organism physiology. Dietary input has a major impact on the stem cell supported ovary in *Drosophila melanogaster* females, appropriately matching reproductive output to nutrient availability. Previous work has shown that inter-organ communication plays a role in modulating the ovarian response to diet. Specifically, amino acid sensing by the adipose tissue remotely controls germline stem cells and their progeny. While we have shown that activation of the amino acid response pathway, a part of the integrated stress response, and mTOR signaling in adipocytes impacts germline stem cell maintenance and ovulation, it is unclear how downstream signaling mediates these responses. Here, using a combination of genetic and cell biological tools, we show that regulation of translation in adult adipocytes impacts the ovarian germline stem cell lineage, from stem cell maintenance to ovulation of mature oocytes. This work strongly suggests that the adipose tissue produces specific factors to control stem cell activity in the ovary and highlights how inter-organ communication underlies organismal physiological responses to diet.

## 1 Introduction

Across a wide range of organisms and tissue types, the proliferative capacity and developmental potential of stem cells support homeostatic maintenance and damage repair. This is especially exemplified in *Drosophila melanogaster*, where physiological factors, like age, sex, hormonal status, and temperature impact adult stem cell activity (Fuller and Spradling, 2007; Ables and Drummond-Barbosa, 2017; Ishibashi et al., 2020; Rodriguez-Fernandez et al., 2020; Finger et al., 2021; Gandara and Drummond-Barbosa, 2022). In particular, dietary input has a major impact on adult stem cell lineages. *D. melanogaster* females fed on a protein-poor diet lay 60-fold fewer eggs compared to females fed a protein-rich diet (Drummond-Barbosa and Spradling, 2001). Studies using the *Drosophila melanogaster* ovary have uncovered many of the cellular and molecular mechanisms that allow adult stem cells and their progeny to respond intrinsically to changes in dietary input.

Adult *D. melanogaster* ovaries are supported by a well characterized germline stem cell (GSC) population. Each ovary consists of approximately 20 individual egg production units called ovarioles (Fig. 1A,B). At the anterior-most tip of each ovariole, the germarium harbors two to three GSCs attached to somatic cap cells, a major component of the stem cell niche (Fig. 1C). GSCs fuel oogenesis, the process of egg production, by dividing asymmetrically giving rise to a GSC and a cystoblast that further undergoes four synchronous divisions with incomplete cytokinesis. This generates a 16-cell germline cyst, which is enveloped by a layer of follicle cells before being pinched off the germarium to form an egg chamber. One cell of the 16-cell germline cyst becomes the oocyte with the remaining 15 nurse cells serving in a support role to the oocyte. The egg chamber progresses through 14 stages to form a mature oocyte ready for ovulation and then fertilization (Fig. 1B) (Giedt and Tootle, 2023). Nutrient status influences multiple steps during the process of oogenesis, such as GSC maintenance and proliferation, germline cyst survival, progression through vitellogenesis, and ovulation (Laws and Drummond-Barbosa, 2017). Moreover, the highly conserved nutrient sensing pathways, insulin/insulin-like growth factor signaling (IIS), mechanistic Target of rapamycin (mTOR) signaling, and AMPK signaling, have been shown to function within ovarian somatic and germ cells to mediate the effects of diet on oogenesis (Laws and Drummond-Barbosa, 2017).

**Figure 1.**
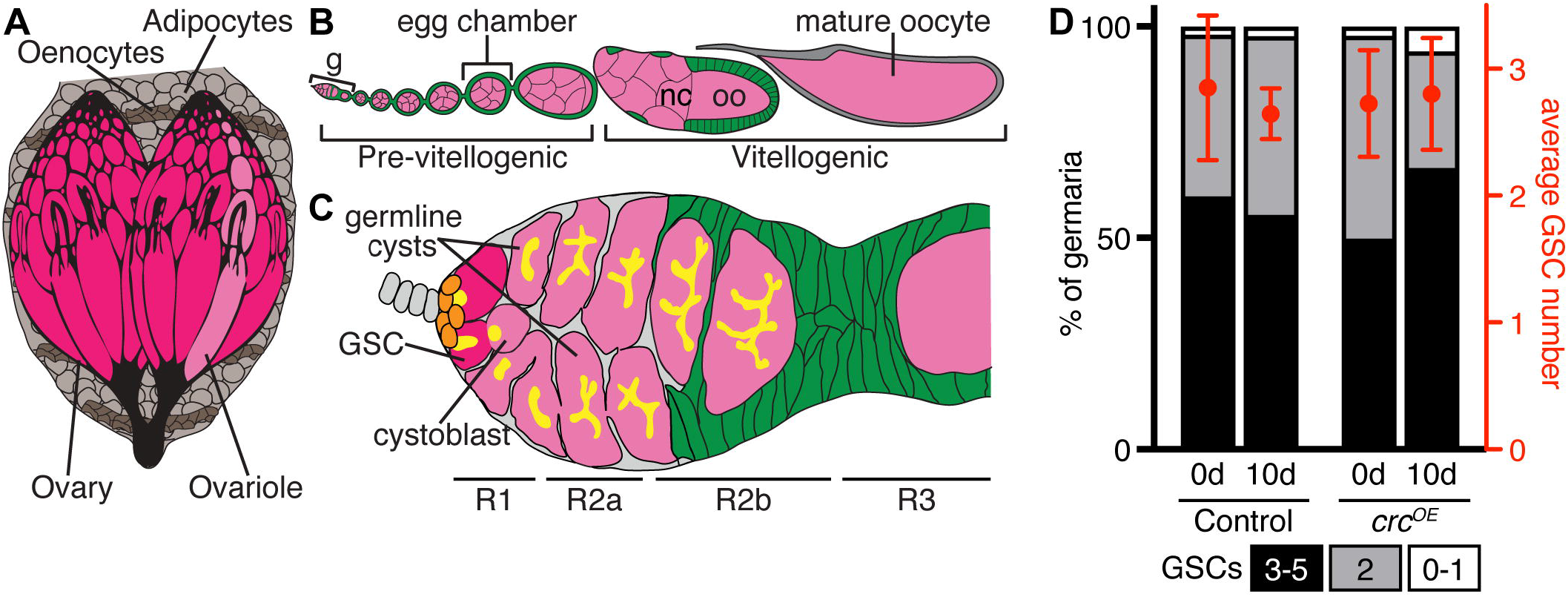
*crc/ATF4* activity in adult adipocytes does not affect germline stem cell maintenance. **(A)** Adipocytes (light brown) and oenocytes (dark brown) make up the adult *Drosophila* fat body which surrounds the paired ovary (pink). **(B)** Adult *D. melanogaster* females have a pair of ovaries that consist of 16-20 ovarioles. Each ovariole contains progressively developing egg chambers. **(C)** The germarium, anterior-most structure of each ovariole, two-three germline stem cells (GSC, magenta) adhere to somatic cap cells (orange). Asymmetric GSC division leads to generation of a cystoblast that undergoes four rounds of synchronous mitotic divisions with incomplete cytokinesis that generates a 16-cell cyst in which germ cells are interconnected by the fusome (yellow). A layer of somatic follicle cells (green) envelops each 16-cell cyst before they exit the germarium to complete the 14 stages of oogenesis that produce a mature oocyte. **(D)** Quantification of the frequencies of germaria with zero-to-one, two, or three-to-five GSCs before and after 10 days of *crc/ATF4* overexpression in adult adipocytes. Average GSC number is shown on the right y-axis (mean ± s.e.m.). Number of germaria analyzed is inside bars.

In addition to ovarian intrinsic nutrient sensing, inter-organ communication, specifically fat-to-ovary, regulates the ovarian response to diet. The *D. melanogaster* fat body, composed of adipocytes and oenocytes (hepatocyte-like cells), is analogous to mammalian adipose tissue with its energy storage and endocrine roles (Arrese and Soulages, 2010). IIS within adult adipocytes, dependent and independent of Akt, controls germline stem cell maintenance, germline cyst survival, and the progression through vitellogenesis (Armstrong and Drummond-Barbosa, 2018). More recently, we have shown that this may be in part mediated by the Ras/MAPK downstream signaling axis (Bradshaw et al., 2024). Regarding sensing of dietary protein status, previous studies have shown that two distinct amino acid sensing pathways function within adult adipocytes to control different steps of oogenesis. The amino acid response (AAR) pathway, part of the integrated stress response, senses limitations in dietary amino acids to reduce global translation and induce expression of ATF4-dependent stress response genes (Murguía and Serrano, 2012). Conversely, mTOR signaling functions during nutrient replete dietary conditions to control protein synthesis, lipid metabolism, and autophagy (Deleyto-Seldas and Efeyan, 2021; Szwed et al., 2021). Interestingly, in *D. melanogaster* adipocytes, AAR pathway activity modulates GSC maintenance while mTOR-mediated signaling promotes ovulation (Armstrong et al., 2014).

Given that the AAR pathway and mTOR both regulate translation, we asked if translation in adipocytes mediates amino acid sensing control of oogenesis. Using genetic tools to knockdown expression of a ribosomal subunit and translation initiation factors, we show that components of the translation machinery are required in adult adipocytes to promote GSC maintenance and ovulation, likely downstream of AAR and mTOR pathway activity. In addition, we find that translation in adipocytes regulate germline cyst survival, likely independent of the AAR and mTOR pathways. Taken together, this work provides evidence that nutrient sensing pathways converge on the molecular process of translation to mediate adipose tissue remote control of oogenesis.

## 2 Results

### 2.1 ATF4-dependent transcription in adipocytes does not influence germline stem cell number

Previous work shows that genetic activation of the AAR pathway in adult adipocytes leads to ovarian GSC loss (Armstrong et al., 2014). Amino acid deprivation activates the AAR pathway, where increased uncharged tRNAs activate the GCN2 kinase which in turn phosphorylates eIF2alpha, ultimately leading to reduced global translation and ATF4-dependent transcription of stress response genes (Murguía and Serrano, 2012). We first asked if ATF4 activity mediates adipocyte control of GSC maintenance by using an adipocyte-specific driver, *tubP-Gal80*^*ts*^*;Lsp2(3*.*1)-Gal4* (Lazareva et al., 2007; Armstrong et al., 2014), to induce overexpression of *cryptocephal* (*crc, ATF4* in *Drosophila*) (Hewes et al., 2000). We find that germaria from flies in which *crc* is overexpressed in adipocytes have average GSC numbers comparable to controls, with similar percentages of germaria containing three-to-five, two, and one-to-zero GSCs (Fig. 1D). Correspondingly, we did not observe a change in cap cell number after 10 days of *crc* overexpression (data not shown). These data suggest that crc/ATF4-dependent transcription does not mediate adipocyte amino acid sensing regulation of GSC number.

### 2.2 Translation machinery is required in adult adipocytes to promote germline stem cell maintenance

To begin addressing if AAR pathway activity suppresses global translation to mediate adipocyte control of GSC maintenance, we asked if reducing intracellular amino acid transport or eukaryotic translation initiation factor 4E1 (*eIF4E1*) in adipocytes affects translation. We used a puromycin incorporation assay to measure new protein synthesis (Deliu et al., 2017) in adipose tissue from flies with adipocyte-specific, RNAi-mediated knockdown of *CG1628*, encodes an amino acid transporter, or *eIF4E1*. Compared to control adipose tissue, we observed greatly reduced puromycin labeling in *Lsp2 > CG1628i* and *Lsp2 > eIF4E1i* fat bodies (Fig. S1). Since reduced amino acid levels activate the AAR pathway which in turn suppresses translation, we predicted that reducing expression of translation machinery components in adipocytes would phenocopy the GSC loss observed when the AAR pathway is experimentally activated (Armstrong et al., 2014). RNAi-mediated knockdown of translation machinery components led to decreased transcript levels of *RpL17, eIF4E1*, and *eEF1β* (Fig. S2). Ten days of adipocyte-specific knockdown of *RpL17*, which encodes a large ribosomal subunit, *eIF4E1*, or *eEF1β*, which encodes eukaryotic translation elongation factor 1 beta, resulted in decreased average GSC number (Fig. 2A), with a higher percentage of germaria containing one or zero GSCs compared to controls (Fig. 2B). Similar to previous observations amino acid transporter knockdown (Armstrong et al., 2014), we find no effect on cap cell number when components of the translation machinery are knocked down in adipocytes (Fig. S3). These data indicate that translation plays a role in mediating the effect of adipocyte AAR pathway activity on GSC maintenance.

**Figure 2.**
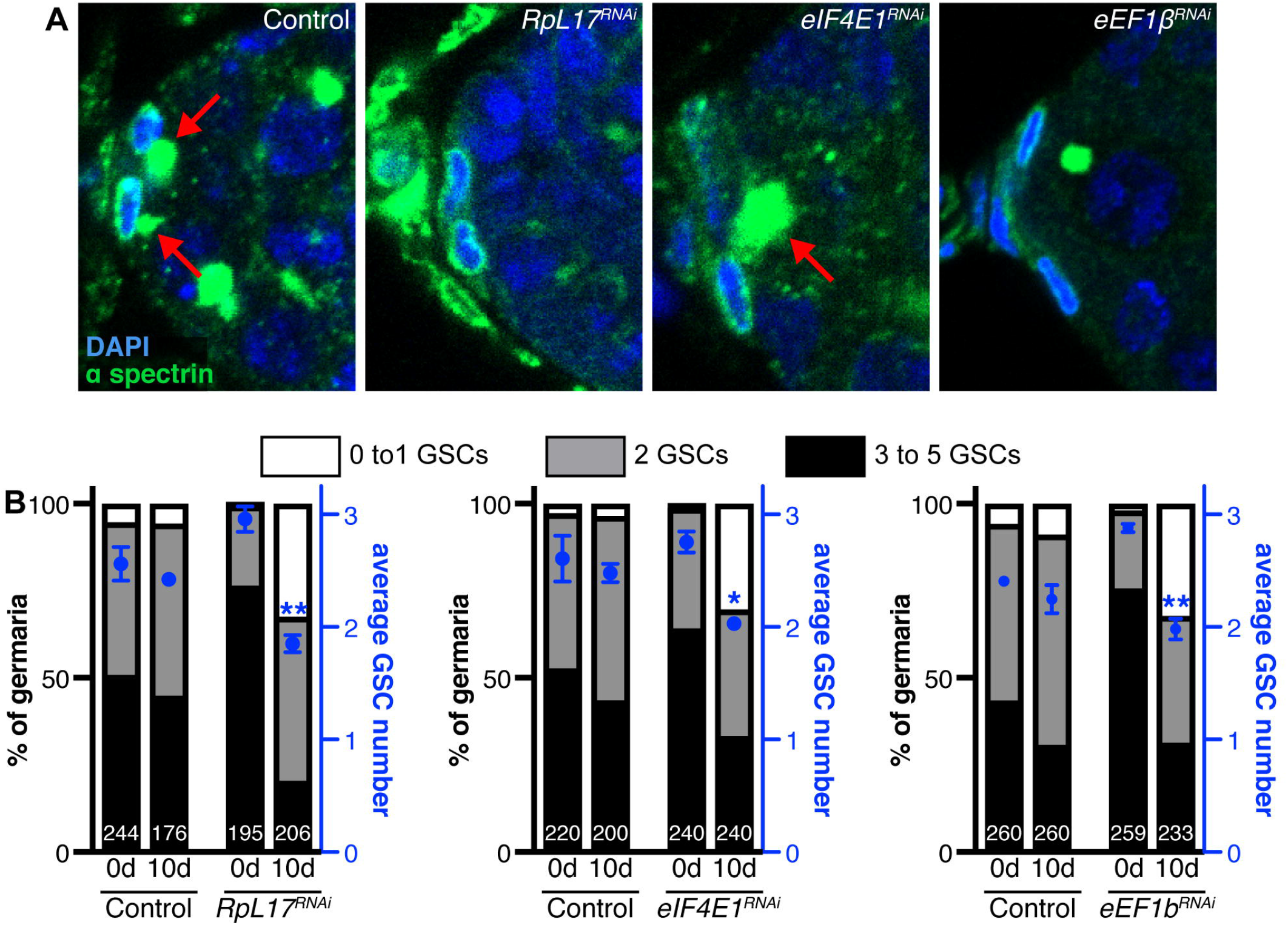
Adipocyte-specific knockdown of translation components leads to GSC loss. **(A)** Representative images of the anterior region of germaria from controls or adipocyte knockdown of *RpL17, eIF4E1*, and *eEF1β* (α-spectrin in green, nuclear outline = cap cells, round/ellipsoid = fusome, arrows; DAPI in blue, nuclei). **(B)** Quantification of the frequencies of germaria with zero-to-one, two, or three-to-five GSCs before and after 10 days of *RpL17, eIF4E1*, or *eIF1β* adipocyte knockdown. Average GSC number is shown on the right y-axis (mean ± s.e.m.). Number of germaria analyzed is inside bars. ** *p* < 0.01, * *p* < 0.05, two-way ANOVA with interaction.

### 2.3 Enhanced expression of translation machinery rescues GSC loss caused by reduced amino acid transport

Next, we asked if enhancing translation could rescue the GSC loss caused by activation of the AAR pathway. Using standard genetic crosses, we generated a fly line containing an RNAi transgene targeting *CG1607*, an amino acid transporter, and an overexpression transgene for *RpL17* (*UAS-CG1607*.*RNAi; UAS-RpL17*.*OE)* to be used in a genetic interaction experiment. As we showed previously, amino acid transporter (*CG1607*) or *RpL17* knockdown in adult adipocytes leads to reduced GSC number (Armstrong et al., 2014)(Fig. 3). However, GSC number was comparable in controls and flies with simultaneous *CG1607* knockdown and *RpL17* overexpression (Fig. 3). This restoration of GSC number provides more direct evidence that the AAR pathway suppresses global translation in adipocytes to modulate GSC number.

**Figure 3.**
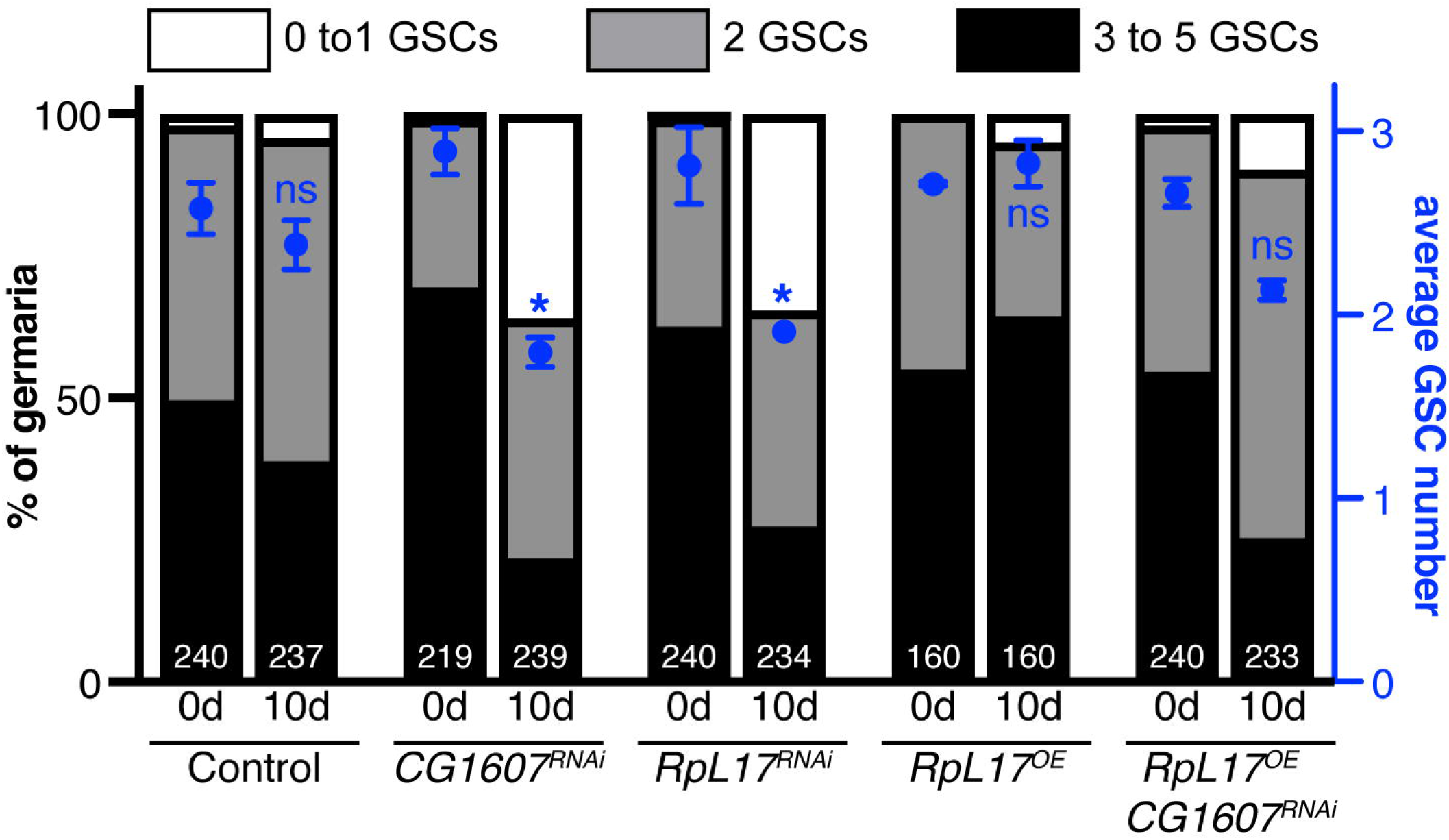
*RpL17* overexpression rescues GSC loss associated with amino acid transporter knockdown in adipocytes. Quantification of the frequencies of germaria with zero-to-one, two, or three-to-five GSCs before and after 10 days of *CG1607* RNAi, *RpL17* RNAi, *RpL17* overexpression, or simultaneous *CG1607* RNAi and *RpL17* overexpression. Average GSC number is shown on the right y-axis (mean ± s.e.m.). Number of germaria analyzed is inside bars. * *p* < 0.05, two-way ANOVA with interaction.

### 2.4 Translation machinery in adult adipocytes promotes germline cyst survival and ovulation

While nutritional regulation of oogenesis is largely influenced by effects on GSC maintenance and proliferation, dietary manipulations impact additional steps of oogenesis. For example, ovaries from female flies fed a protein-poor diet have delayed egg chamber development and blocked vitellogenesis (Drummond-Barbosa and Spradling, 2001), similar to ovaries from female flies fed a high-sugar diet (Nunes and Drummond-Barbosa, 2023). Like GSCs, early and late germline development respond to nutrient sensing pathway activity in the fat body. Previous studies show that Akt-dependent IIS and Ras/MAPK axis signaling are required in adult adipocytes to support germline cyst survival (Armstrong and Drummond-Barbosa, 2018; Bradshaw et al., 2024). Additionally, adipocyte mTOR signaling promotes ovulation (Armstrong et al., 2014).

Therefore, we first asked if adipose tissue control of germline cyst survival could be mediated by adipocyte translation. We knocked down *RpL17, eEF1β*, *eIF4E1*, and *eIF2α* in adult adipocytes and quantified germline cyst death in germaria as evidenced by Dcp-1 immunoreactivity (Fig. 4A). When measuring cell death in whole germaria from females with adipocyte-specific knockdown of translation components, we observed a higher percentage of Dcp-1 labeling compared to controls, with statistically significant increases for knockdown of *RpL17* and *eIF2α* (Fig. 4B). Moreover, region one of the germarium, which houses mitotically dividing germ cells (de Cuevas and Spradling, 1998), showed large, statistically significant increases in Dcp-1 immunoreactivity for all knocked down translation components compared to controls (Fig. 4B). Next, we asked if adipose tissue control of ovulation could be mediated by adipocyte translation. In most ovaries from control females, each ovariole contains one or no mature stage 14 oocytes (Fig. 5A) with some ovaries containing at least one ovariole with more than one stage 14 oocyte (we refer to this as blocked ovulation) (Fig. 5B, an average of 37.5%). Conversely, most ovaries from females in which *RpL17* (an average of 76%) or *eIF2α* (an average of 86%) has been knocked down in adipocytes show the blocked ovulation phenotype (Fig. 5). Taken together, these data suggest that translation in adipocytes regulates germline cyst survival and ovulation.

**Figure 4.**
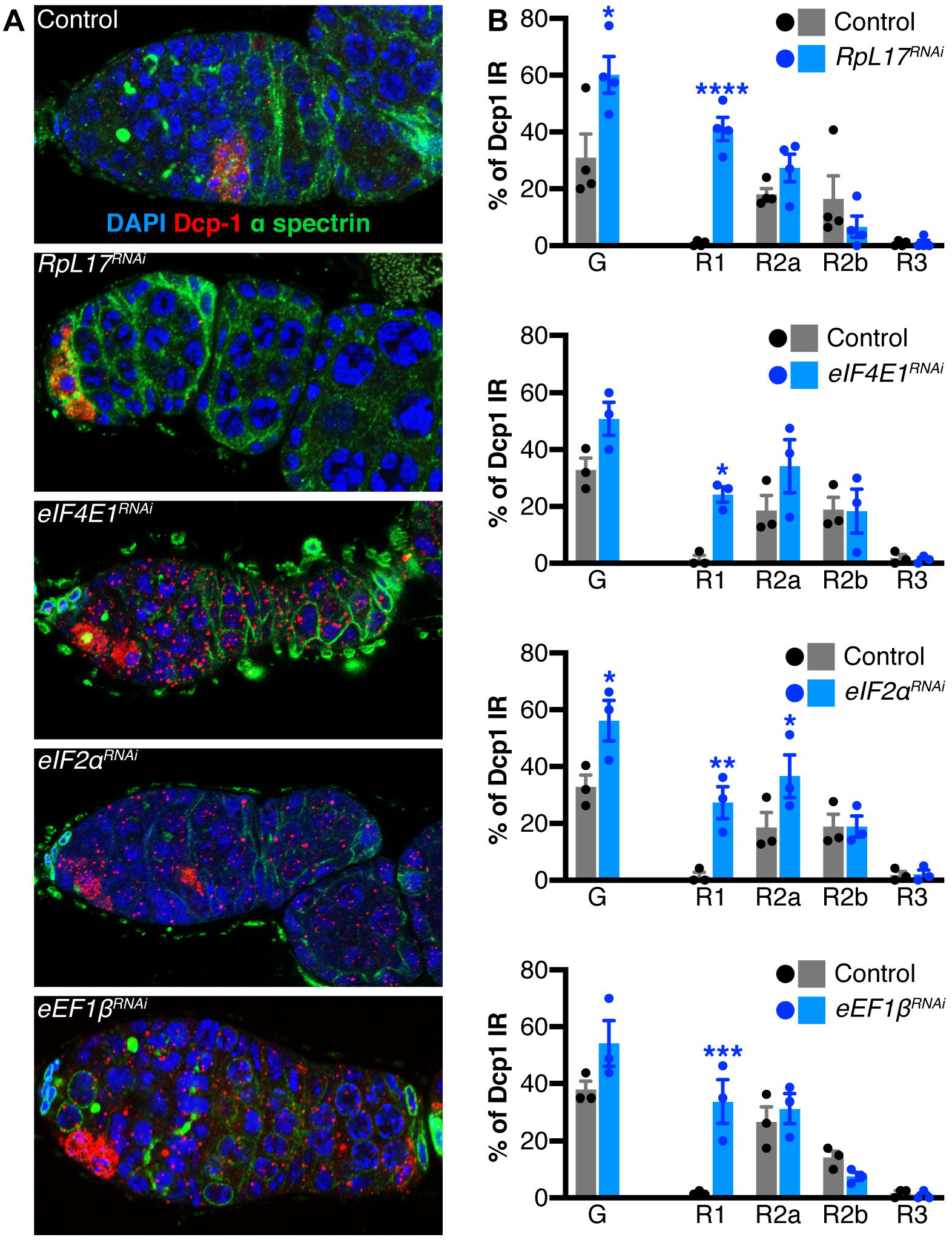
Adipocyte-specific knockdown of translation components leads to increased germline cyst death. **(A)** Representative germarium images from controls or adipocyte knockdown of *RpL17, eIF4E1, eIF2α*, and *eEF1β* (α-spectrin in green, cellular membranes; Dcp-1 in red, dying cysts; DAPI in blue, nuclei). **(B)** Percentage of Dcp-1 immunoreactivity in whole germaria (G) and by germarium region (R1, R2a, R2b, and R3) from female flies with adult adipocyte-specific knock down of *RpL17, eIF4E1, eIF2α*, and *eEF1β*. Number of germaria are XYZ that were analyzed over three biological replicates (shown as individual data points on top of bars). **** *p* < 0.0001, *** *p*, 0.001, ** *p* < 0.01, * *p* < 0.05, Student’s two-tailed *t*-test (experimental group compared to its corresponding control).

**Figure 5.**
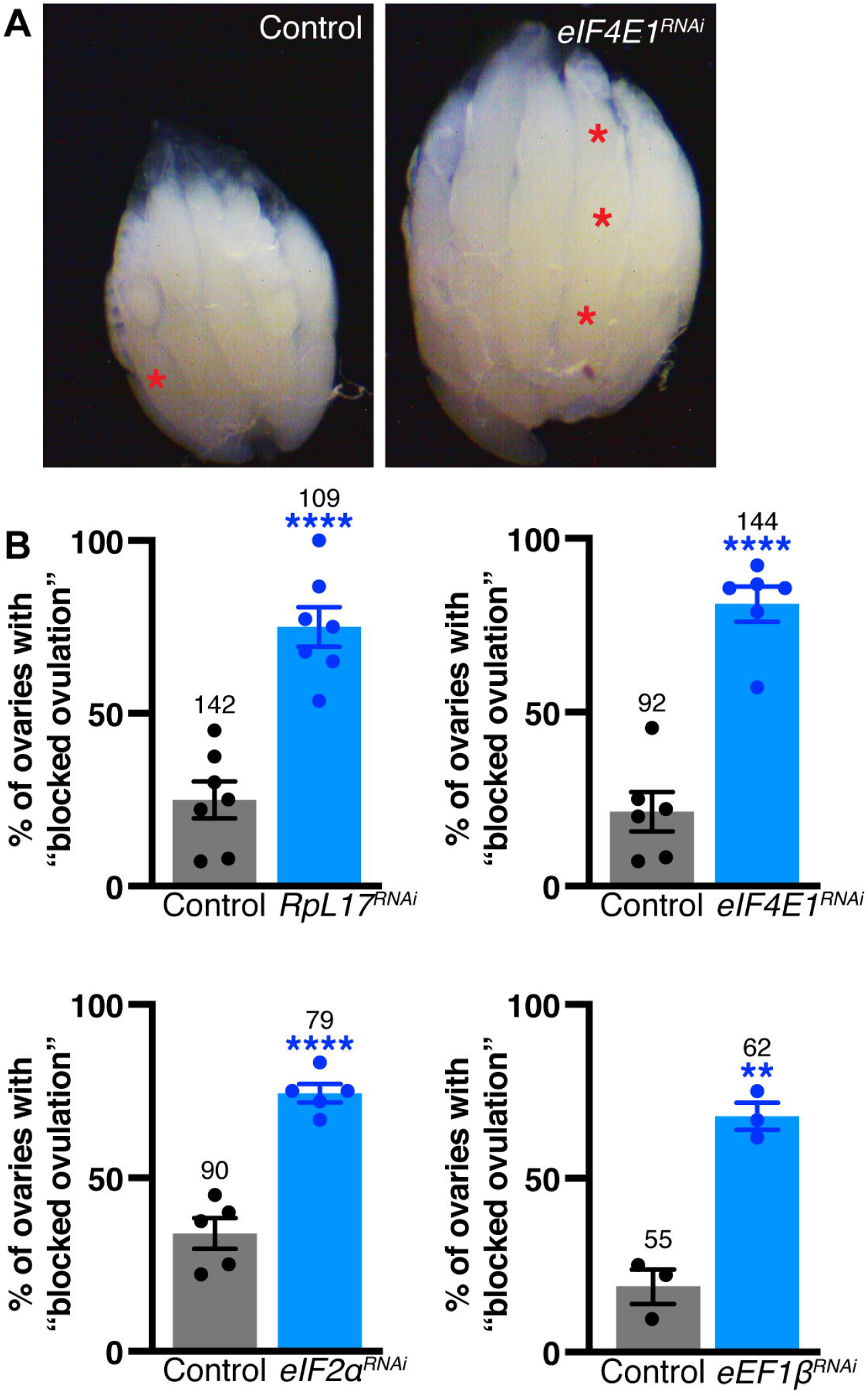
Adipocyte-specific knockdown of translation components leads to blocked ovulation. **(A)** Representative ovaries from controls and females with adipocyte-specific knockdown of eIF4E1. Asterisks mark stage 14 oocytes. **(B)** Percentage of ovaries containing at least one ovariole that retains more than one oocyte at 10 days of adult adipocyte-specific knock down of *RpL17, e1F4E1, eIF2α*, or *eEF1β*. Number of ovaries analyzed over three-six biological replicates (indicated as individual data points) is shown inside bars.

## 3 Discussion

In addition to individual, direct tissue responses, multi-organ communication allows for a coordinated, whole-organism response to changes in dietary input. This is particularly important in organisms for which food availability can vary greatly with respect to amount, quality, and frequency and for tissues that are highly sensitive to nutritional input, like the *Drosophila melanogaster* ovary. It is well-established that the *Drosophila* adipose tissue controls oogenesis (Armstrong et al., 2014; Matsuoka et al., 2017; Armstrong and Drummond-Barbosa, 2018; Weaver and Drummond-Barbosa, 2018, 2019; Bradshaw et al., 2024). Moreover, activity of nutrient sensing pathways, like IIS, mTOR, and AAR, in adipocytes are known to regulate distinct aspects of oogenesis (Armstrong et al., 2014; Armstrong and Drummond-Barbosa, 2018; Bradshaw et al., 2024). In this study, we show that translation within adipocytes mediates the ovarian response to nutrient sensing by the fat body. Specifically, we find that translation components are required within adipocytes to support GSC maintenance, germline cyst survival, and ovulation of mature oocytes. Together with previous studies showing that the PI3K/Akt and Ras/MAPK downstream axes in adipocytes control different stages of oogenesis, the work described here indicates that activity of distinct signaling pathways in adipocytes converge on translation regulation to produce adipocyte-derived factors that relay information to the ovary.

Convergence on translation regulation may underlie the ability of multiple nutrient-sensing signaling pathways to control oogenesis, with the production of individual proteins corresponding to specific steps in the process. The Ras/MAPK signaling axis activates p90 ribosomal S6 kinases (RSKs) which phosphorylate rpS6, a protein component of the small ribosomal subunit, independent of mTOR to regulate translation initiation (Roux et al., 2007). AAR pathway-dependent activation of the GCN2 kinase leads to suppression of global translation (Wek et al., 2023). Activity of the PI3K/Akt signaling axis inhibits the mTOR inhibitory complex, TSC1/TSC2, thus allowing mTOR to promote translation by phosphorylation of 4E-BP and S6K (Yang et al., 2022). Given that these pathways have been shown to function in adult adipocytes to control GSC maintenance, germline cyst survival, vitellogenesis, and ovulation (Armstrong et al., 2014; Armstrong and Drummond-Barbosa, 2018; Bradshaw et al., 2024), they could be controlling the production of protein factors that communicate directly or indirectly to the ovary. Interestingly, Ras/MAPK, PI3K/Akt, and AAR pathway signaling have been shown to regulate mTOR (Cargnello and Roux, 2011; Hu and Guo, 2021; Yang et al., 2022), thus suggesting that mTOR-dependent translation might be the primary mediator of fat-to-ovary communication about nutrient input. However, InR activity, but not mTOR signaling, in adipocytes is required for GSC maintenance (Armstrong et al., 2014; Armstrong and Drummond-Barbosa, 2018) indicating that at least PI3K/Akt- and mTOR-dependent translation are separable. Future studies employing the robust *Drosophila melanogaster* genetic and molecular toolkit will decipher how differential inputs on translation regulation in the adipose, and other tissues, mediate distinct effects on oogenesis.

There is clear evidence that the *D. melanogaster* adipose tissue communicates information about dietary protein to the ovary in adult flies, yet the identity of protein factors that mediate this inter-organ communication and which nutrient-sensing pathways they respond to remain unknown. Hundreds of fat body proteins show rapid changes in diet-dependent expression when adult females are switched from a protein-rich to a protein-poor diet (Matsuoka et al., 2017). Additionally, the adult fat body secretes over 1,500 newly synthesized proteins into the hemolymph (Droujinine et al., 2021). When cross-referencing these two datasets, we find that several proteins that are down-regulated on a protein-poor diet are also known to be secreted from the fat body and detected in the hemolymph. Moreover, Tain and colleagues identified 256 *D. melanogaster* fat body proteins that are differentially expressed when IIS is perturbed (Tain et al., 2017). Translation of nearly 3,000 proteins is affected in mammalian cells treated with an inhibitor of mTOR-dependent translation (Klann et al., 2020). We postulate that these proteins are candidate factors whose expression might be regulated by adipocyte IIS, mTOR, and AAR pathways. Therefore, future studies in which expression of individual putative fat body-derived factors are knocked down in adult adipocytes will reveal specific proteins that regulate steps of oogenensis that are sensitive to dietary amino acids.

## 4 Materials and Methods

### 4.1 *Drosophila* strains and experimental conditions

Fly stocks were maintained at 22-25°C on medium containing cornmeal, molasses yeast, and agar. For protein rich dietary conditions, standard medium was supplemented with wet yeast paste to induce robust egg production. For adipocyte-specific manipulation of gene expression, the *3*.*1Lsp2-Gal4* driver line (Lazareva et al., 2007) was used. For temporal control of gene expression, the *3*.*1Lsp2-Gal4* driver was combined with a temperature-sensitive *Gal80* under the control of a ubiquitous promoter (*tubP-Gal80*^*ts*^)(McGuire et al., 2003; Armstrong et al., 2014). For knockdown of translation components and amino acid transporters, *UAS-RNAi* lines were procured from the Vienna Drosophila RNAi Stock Center (VDRC, http://stockcenter.vdrc.at) or the Transgenic RNAi Project (TRiP, http://www.flyrnai.org) collections. *UAS-RNAi* lines used are listed in supplementary material Table S1. The *UAS*-*CG1607* RNAi, *UAS* -*RpL17* overexpression line was generated by standard crosses for this publication. Other genetic elements used are described in FlyBase (http://www.flybase.org).

### 4.2 Adult adipocyte-specific genetic manipulations

Un-mated females from the *tub-Gal80*^*ts*^; *3*.*1Lsp2-Gal4/TM6b* driver line were mated with *UAS-transgene* males. *UAS-GFP*^*dsRNA*^ was used as a control. Crosses were maintained at 18°C, the permissive temperature for *Gal80*^*ts*^, for approximately three weeks to suppress RNAi induction during larval development. Upon eclosion of progeny, virgin females of the desired genotype (*tub-Gal80ts; 3*.*1Lsp2-Gal4 > UAS-transgene* or *tub-Gal80ts; 3*.*1Lsp2-Gal4 > UAS-transgene1; UAS-transgene2*) were collected and mated with *w*^*1118*^ males. After four days at 18°C, flies were switched to 29°C, the restrictive temperature for Gal80^ts^, and fed fresh food daily to induce transgene expression for 10 days prior to dissection. Ovary samples obtained after four days at 18°C, prior to transgene expression, is defined as the 0-day (0d) time point. Ovary samples obtained after four days at 18°C followed by 10 days at 29°C is defined as the 10-day time point (10d).

### 4.3 Whole-mount ovary immunostaining and fluorescence microscopy

Ovaries were dissected and ovarioles partially teased apart in Grace’s medium (Caisson Labs) and fixed in 5.3% paraformaldehyde (Electron Microscopy Sciences) for 13 minutes at room temperature. Ovary samples were rinsed three times them washed three times in 0.1% Triton X-100 (VWR Life Sciences) in 1X phosphate buffered saline (PBT), and then blocked in 5% bovine serum albumin (BSA, US Biological Life Sciences) and 5% normal goat serum (NGS, MP Biomedicals, LLC) in PBT (blocking solution) overnight at 4°C or room temperature on a nutator. Ovaries were incubated overnight at room temperature in the following primary antibodies, mouse anti-α-spectrin (1:50, Developmental Studies Hybridoma Bank [DSHB]), mouse anti-Lamin C (LamC, 1:50, DSHB), and rabbit anti-cleaved *Drosophila* Dcp-1 (1:200, Cell Signaling Technology). The ovaries were then washed three times for 15 minutes in 0.1% PBT followed by incubation with Alexa Fluor 488- and 568-goat species-specific secondary antibodies (1:250, Invitrogen) for two hours at room temperature and protected from light. After three, 15 minute 0.1% PBT washes, ovary samples were mounted in Vectashield containing 4′,6-diamidino-2-phenylindole (DAPI, Vector Laboratories). Fluorescence images of germaria were acquired using the 63X oil objective on a Zeiss LSM 800 confocal microscope equipped with 2.6ZEN software.

### 4.4 Ovarian analysis

Cap cells were identified based on enriched Lamin C staining, nuclear morphology, and anterior germarium location. GSCs were identified based on their juxtaposition to cap cells and fusome morphology (de Cuevas and Spradling, 1998) revealed by α-spectrin staining. Cap cell and GSC numbers were counted for each germarium at 0d and 10d of transgene expression from at least three independent experiments. For statistical analysis for differences in GSC loss over time two-way ANOVA with interaction (GraphPad Prism 8) was performed, as described (Armstrong et al., 2014). Germline death was identified by Dcp-1 immunoreactivity and scored as the number of germaria with at least one dying cyst (using fusome morphology) at 10d of transgene expression from at least three independent experiments. The percentage of germline cyst death was calculated as the number of Dcp-1-positive germaria divided by the total number of germaria analyzed, multiplied by 100. Data for germline cyst death for the whole germarium, irrespective of regions, was subjected to unpaired Student’s *t*-tests while one-way ANOVA was used to compare germline cyst death per region (GraphPad Prism 8). For ovulation analyses, whole ovaries dissected at 10d were examined using a Zeiss Stemi 508 stereomicroscope. Ovaries containing more than one mature oocyte were categorized as having blocked ovulation. Bright-field images of whole ovaries were captured on a Zeiss Stemi 305 stereomicroscope with the Labscope imaging app and data was analyzed using one-way ANOVA (GraphPad Prism 8).

### 4.5 RNA isolation and RT-PCR

Abdominal carcasses, i.e., abdomens lacking internal organs, from 7-10 females per genotype were hand dissected in DNA/RNA Shield (Zymo Research) after 10 days of transgene induction. After scraping fat body tissue from abdominal carcasses, RNA was extracted using the Quick-RNA Miniprep Kit (Zymo Research) according to the manufacturer’s instructions. Using 1μg of RNA, cDNA synthesis was carried out using the Verso cDNA Synthesis Kit (Thermo Fisher Scientific) according to the manufacturer’s instructions. For each primer pair PCR was performed for the control and corresponding RNAi or overexpression samples using EconoTaq PLUS Green (Lucigen) on an iCycler Thermocycler (Bio-Rad). Primers used are listed in supplemental material Table S2 (*rp49* primers were used as a control). Band intensities were quantified using FIJI software by subtracting background pixels from band pixels and then normalizing to the corresponding housekeeping gene band (rp49 or Tub alpha). Controls were set to one and the experimental sample intensities were determined relative to the control.

### 4.6 Puromycin incorporation assay and western blotting

On day 10 of transgene expression, fat body from 10 to 15 female flies were hand dissected in Grace’s medium such that the fat body tissue was still attached to the abdominal carcass. Abdominal carcasses were incubated in 50 μg/ml puromycin (VWR) diluted in Grace’s medium for one hour at room temperature on a nutator to allow proper penetration. Following two, five minute washes in 1X PBS, abdominal carcasses were homogenized in RIPA buffer using a motorized pestle and then passed through using 22 gauze syringe. Homogenized samples were incubated on ice for 30 minutes in RIPA buffer followed by centrifugation for 15 min at 4°C at 15000rpm. The supernatant was carefully collected without disturbing the pellet and analyzed using western blot. Total protein quantification was carried out using a BCA Protein assay kit (Thermo Fisher Scientific). 15 μg of total protein was used for SDS-PAGE. Proteins were transferred onto a nitrocellulose membrane, which was then blocked with 5% skim milk and incubated with mouse monoclonal anti-puromycin antibody (Millipore; 1:1000). Membranes were later stripped of antibodies and probed for anti alpha-tubulin (DHSB; 1:1000). For secondary antibody labeling, membranes that had been washed three times 20 minutes in TBS with 0.1% Tween 20 at room temperature were incubated with HRP-conjugated goat anti-mouse antibody (Cell Signaling Technology; 1:5000). The chemiluminescent signal was developed using the Clarity Western ECL substrate (Bio-Rad) and captured using the Image Lab software (Bio-Rad). Using ImageJ, puromycin densitometry was quantified by subtracting the background, measuring signal intensity, and normalizing to tubulin control bands.

## 6 Author Contributions

S.S. and A.R.A. designed the experiments and interpreted the data as well as wrote the manuscript. S.S. performed all experiments.

## 8 Acknowledgments

We thank the Bloomington Drosophila Stock Center and the TRiP at Harvard Medical School for fly stocks. The monoclonal antibodies developed by P.A. Fisher, D. Branton, R. Dubreuil, and T. Uemura were obtained from the Developmental Studies Hybridoma Bank, created by the NICHD of the NIH and maintained at The University of Iowa, Department of Biology, Iowa City, IA 52242.

