## Supplemental Figures for "Translation components in adult *Drosophila melanogaster* adipocytes regulate the ovarian germline stem cell lineage"

**Sahu and Armstrong – Supplemental Figures**

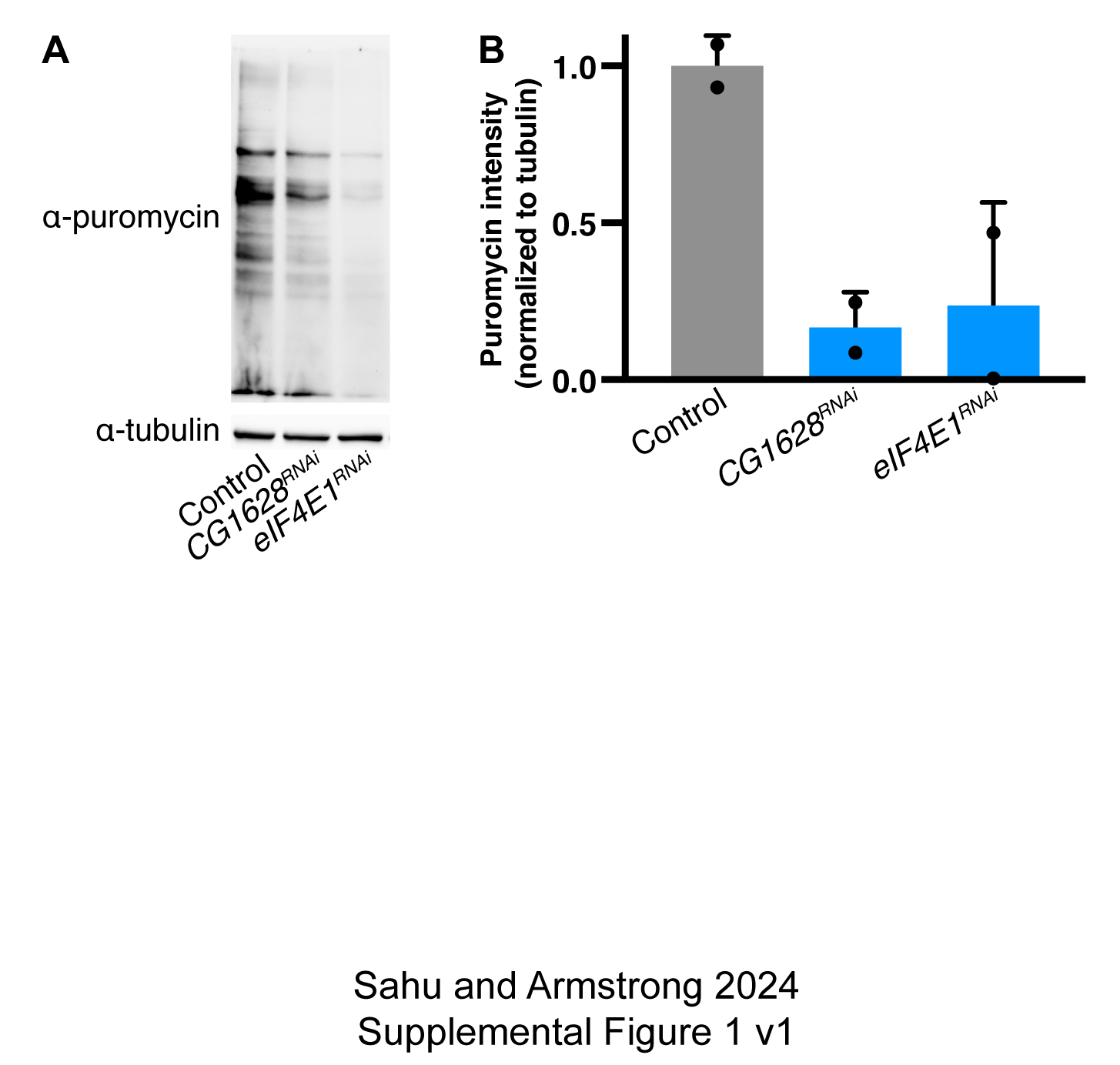

**Supplemental Figure 1**: (A) Adipocyte-specific knock down of *CG1628* or *eIF4E1* leads to reduced puromycin incorporation in adipose tissue. (B) Quantification of two biological replicates, indicated by individual data points.

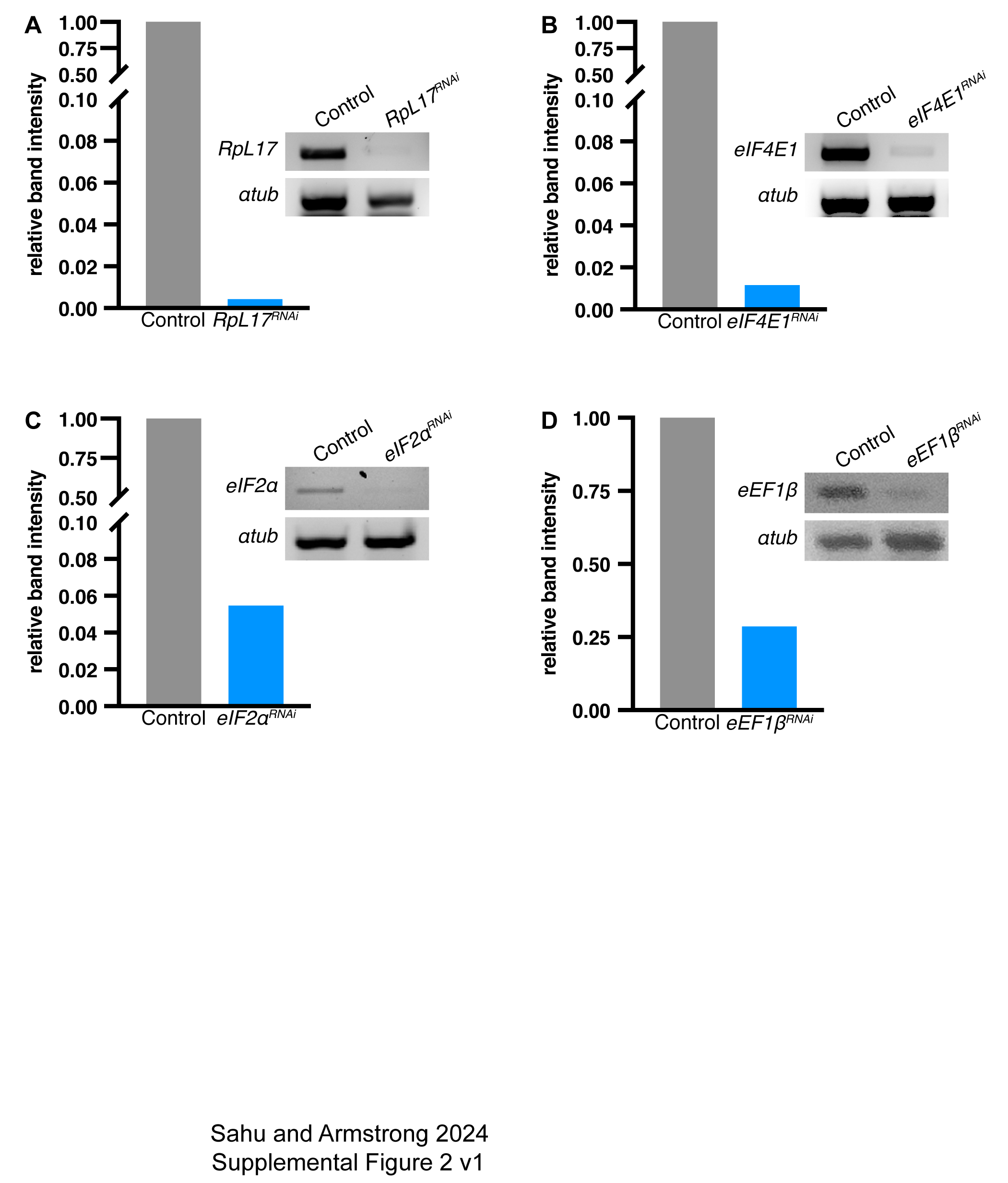

**Supplemental Figure 2**: RT-PCR analysis of mRNA expression in fat bodies with adipocyte-specific, RNAi-mediated knockdown of *RpL17 (A)*, *eIF4E1 (B)*, *eIF2α* (C), and *eEF1β* (D). *α-tubulin* levels for each sample are shown below each gel. Quantification of knockdown efficiency relative to controls (*GFP* RNAi) and normalized to *Rp49* levels are shown in the graphs.

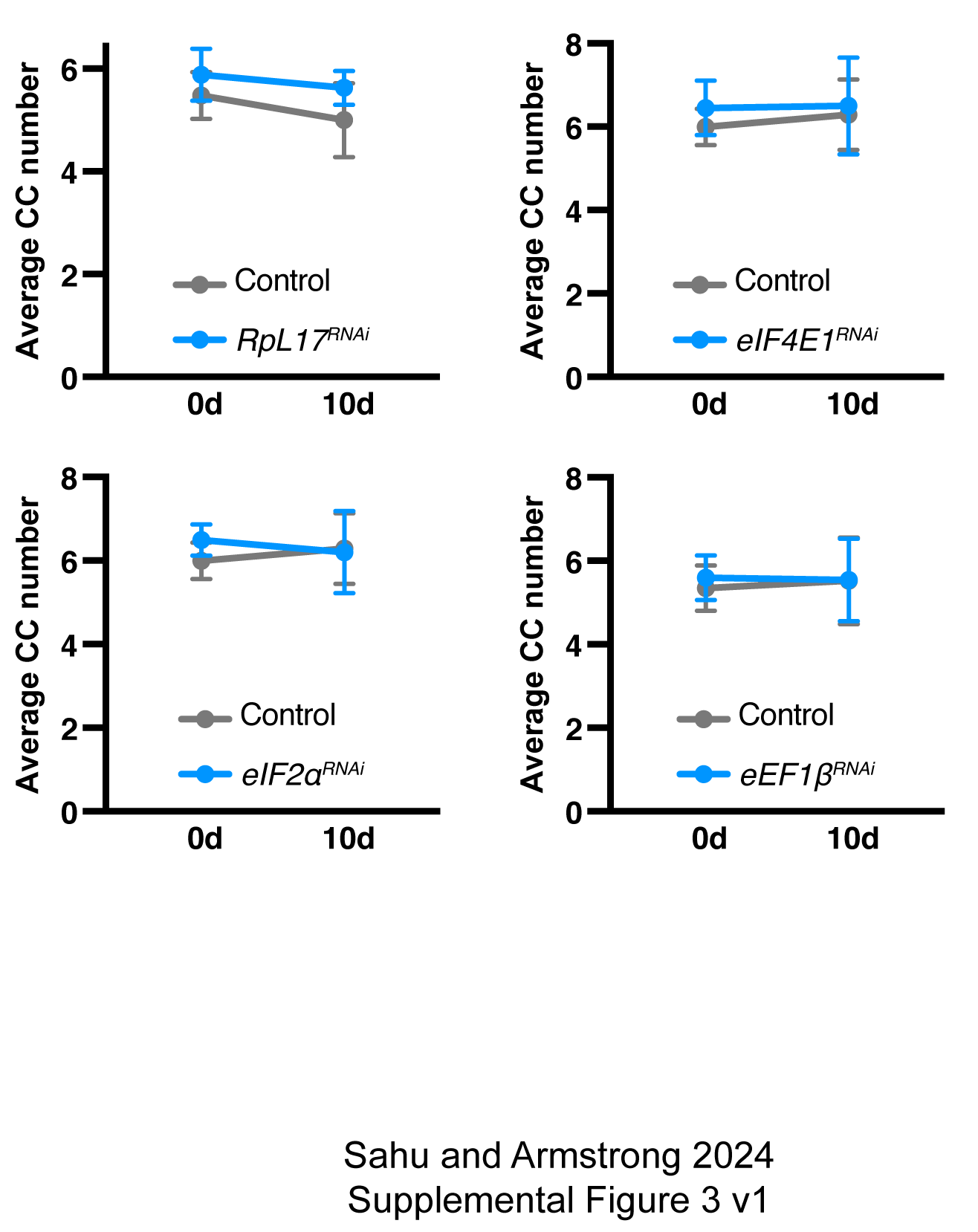

**Supplemental Figure 3**: Adipocyte-specific knockdown of *RpL17*, *eIF4E1*, *eIF2α*, and *eEF1β* do not impact cap cell number. *α-tubulin* levels for each sample are shown below each gel. Means ± s.e.m. are shown for three biological replicates.
