## Supplemental Table 1 for "Translation components in adult *Drosophila melanogaster* adipocytes regulate the ovarian germline stem cell lineage"

**Table S1.** UAS-RNAi lines used in this study.

|  | **Gene** | ***UAS-dsRNA* line** | **Source** |
| --- | --- | --- | --- |
| controls | GFP | *P{UAS-GFP.dsRNA.R}143* | BDSC #9331 |
|  | GFP | *y1 w*; P{UAS-GFP::lacZ.nls}30.1* | BDSC #6452 |
| Amino acid transporter | *CG1607* | *P{GD4651}v14925* | VDRC v14925 |
| Translation machinery | *RpL17* | *w1118; P{GD10434}v41777* | VDRC v41777 |
|  | *RpL17* | *M{UAS-RpL17.ORF.3xHA}ZH-86Fb* | BDSC #F000761 |
|  | *eEf 1* | *w1118; P{GD11894}v45571* | VDRC v45571 |
|  | *eIf4E1* | *w1118; P{GD1432}v7800* | VDRC v7800 |
|  | *eIF2* | *w1118; P{GD1430}v7799* | VDRC v7799 |
|  | *crc* | *yw; UAS-crc[14]/CyO,y+* | Hewes et al., 2000 |
