## Supplemental Table 2 for "Translation components in adult *Drosophila melanogaster* adipocytes regulate the ovarian germline stem cell lineage"

**Table S2.** Primer sequences used in this study.

| Gene | Forward | Reverse |
| --- | --- | --- |
| *eEF1β* | 5’-CATCAGCGGATATACTCCCA-3’ | 5’-AATAGATCCACATCGTCGTC-3’ |
| *eIF1A* | 5’-GACAAAATTTGAAGTCGTTGAG-3’ | 5’-GAATGTGACTGTCTCGTTGA-3’ |
| *eIF2α* | 5’-CACCGCCACTCTATGTAATG-3’ | 5’-ATCTCGCTGTTTTCACTTTT-3’ |
| *eIF4E1* | 5’- ATTAAGCCCAGTAACCTACG-3’ | 5’- CATGGGACGAATGTTCTTCTT - 3’ |
| *MetRs* | 5’-GGTCAGGATTCTAGCTTCAG-3’ | 5’-TAACAAGTGGTTTCTCAGCA-3’ |
| *RpL17* | 5’-TTCCACGTTTCCGATATTGA-3’ | 5’-ACGACATGTAGGGATTGATG-3’ |
| *crc* | 5’- GAGTTGAGTAATGTGCGTTG-3’ | 5’- GATGGCGCAAGAGTCAAT-3’ |
